# Hundreds of motif pairs may facilitate enhancer-promoter interactions

**DOI:** 10.1101/2020.12.29.424656

**Authors:** Saidi Wang, Haiyan Hu, Xiaoman Li

## Abstract

Previous studies have shown that pairs of interacting transcription factors (TFs) bind to enhancers and promoters and contribute to their physical interactions. However, to date, we have limited knowledge about these interacting TF pairs. To identify such TF pairs, we systematically studied the co-occurrence of TF-binding motifs in interacting enhancer-promoter (EP) pairs in seven human cell lines. We discovered hundreds of motif pairs that significantly co-occur in enhancers and promoters of interacting EP pairs. We demonstrated that these motif pairs are biologically meaningful and significantly enriched with motif pairs of known interacting TF pairs. We also showed that the identified motif pairs facilitated the discovery of the interacting EP pairs. The predicted motifs and motif pairs are available at http://www.cs.ucf.edu/~xiaoman/ET/EPmotif/.

## Introduction

Identifying enhancer-promoter (EP) interactions is important for the understanding of gene transcriptional regulation [1]. Enhancers are short genomic regions that can strengthen their target genes’ transcriptional levels independent of their distance and orientation to the target genes [2]. They are in general several hundred base pairs (bps) long, can be hundreds to thousands of bps away from their target genes, and can be in the upstream or downstream of the target genes or in introns. By interacting with promoters of their target genes, enhancers strengthen target genes’ transcription and modulate their condition-specific expression [2–4].

Many studies have attempted to identify EP interactions. Experimental approaches based on chromatin conformation capture techniques and their extensions have identified many EP interactions across several cell lines, cell types and tissues [1, 5–11]. These experimental approaches nurtured our rudimentary understanding of EP interactions. However, they are either time-consuming or still costly because of the large number of EP interactions under an experimental condition and the required high-sequencing depth to comprehensively identify them on the genome-scale [1, 12]. Computational methods for EP interaction predictions are thus indispensable. These methods usually consider the distance, conservation, correlated activity between enhancers and promoters, etc., to identify EP interactions [13–22]. Although having shown success, they have a suboptimal performance on discovering EP interactions, especially condition-specific EP interactions [17, 23–25]. It is thus necessary to further investigate the characteristics of EP interactions, which may significantly facilitate the improvement of the accuracy of the existing methods.

Several studies pointed out a new venue to explore the characteristics of EP interactions, which suggested that the interaction of transcription factors (TFs) that bind an enhancer and TFs that bind a promoter of an EP pair may contribute to the interaction of this EP pair [2, 12, 20, 26–30]. For instance, it is well known that the TF and structural protein CTCF binds to a fraction of enhancers and promoters, which facilitates the physical interaction of enhancers and promoters in these EP pairs [31]. Another example, the ubiquitous TF YY1, binds to enhancers and promoters and contributes to EP interactions as well [32]. It is thus promising to systematically study the potential interactions of TFs that bind to enhancers and promoters and understand how such interactions may lead to the interaction of EP pairs. A computational study integrated chromatin immunoprecipitation followed by massive parallel sequencing (ChIP-seq) data and Hi-C data in two cell lines and predicted 565 interactions of DNA-binding proteins, including TFs [12]. This study was encouraging while limited with a small number of TFs in only two cell lines. To date, we still lack a clear view of the interaction of which TF pairs may render the specificity of the interaction of the enhancer and the promoter in an EP pair.

To address this question, we systematically investigated the co-occurrence of potential TF binding motifs in enhancers and their corresponding interacting promoters (Material and Methods). A motif is a TF binding pattern, which is often represented by a position weight matrix [33, 34]. We identified 114 non-redundant motifs in interacting EP pairs that represented the binding patterns of potential TFs. We also identified 423 motif pairs that significantly cooccurred in interacting EP pairs. Interestingly, on average, more than 62% of these motif pairs in a cell line were shared across cell lines and were able to help to distinguish true interacting EP pairs from false ones. Our study provides a comprehensive list of motif pairs that may contribute to EP physical interactions and facilitate their predictions, which also generates useful hypotheses for experimental validation.

## Material and Methods

### Positive and negative EP pairs

We downloaded the Hi-C contact matrices in the following seven cell lines: GM12878, HMEC, HUVEC, IMR90, K562, KBM7 and NHEK, which were normalized with the Knight and Ruiz normalization vectors by Rao et al. [1]. We claimed that two genomic regions interacted in a cell line (except GM12878) if the corresponding entry in the normalized contact matrix of this cell line was larger than 30. We chose the cutoff 30 because the interacting regions defined by this cutoff would include almost all pairs of interacting regions defined in IMR90 and K562 by independent studies [6, 7]. In GM12878, we used the cutoff 150 instead of 30 since GM12878 had a much larger sequencing depth than other cell lines, and this cutoff resulted in a similar number of pairs of interacting regions to that in other cell lines [1]. In this way, we had positive pairs of interacting regions. Note that we could use looplists defined by Rao et al. as positive pairs of interacting regions [1]. However, the number of looplists was small, which resulted in an even smaller number of positive EP pairs that could not be used to discover interacting TF pairs below. We thus considered the positive pairs of interacting regions defined by the above cutoffs, the majority of which are likely to be reliable because of the large number of supporting reads [1].

To obtain positive EP pairs in a cell line, we overlapped the above positive pairs of genomic regions with the corresponding “active” enhancers and “active” promoters (Fig. 1A). An active enhancer was one of the 32284 enhancers defined by FANTOM [35] that overlapped with the H3K27ac ChIP-seq peaks defined by ENCODE [36] in the corresponding cell line. To our knowledge, FANTOM enhancers were the largest collection of mammalian enhancers with direct experimental evidence. With the transcription start sites (TSSs) defined in GENCODE, we defined 57820 promoters, each of which was the genomic region from the upstream 1000 bps to the downstream of 100 bps the TSS of a GENCODE gene. An active promoter was then defined with these GENCODE promoters and the ENCODE RNA-seq data similarly as previously [19, 24]. In this way, every positive EP pair had an active enhancer overlapping with one genomic region and an active promoter overlapping with the other genomic region of a positive pair of genomic regions, and the distance between the enhancer and the promoter was within 2.5 kilobase pairs to 2 megabase pairs (Supplementary S1). The majority of the positive EP pairs were likely to be true positives, despite false positives and false negatives.

**Fig. 1.**
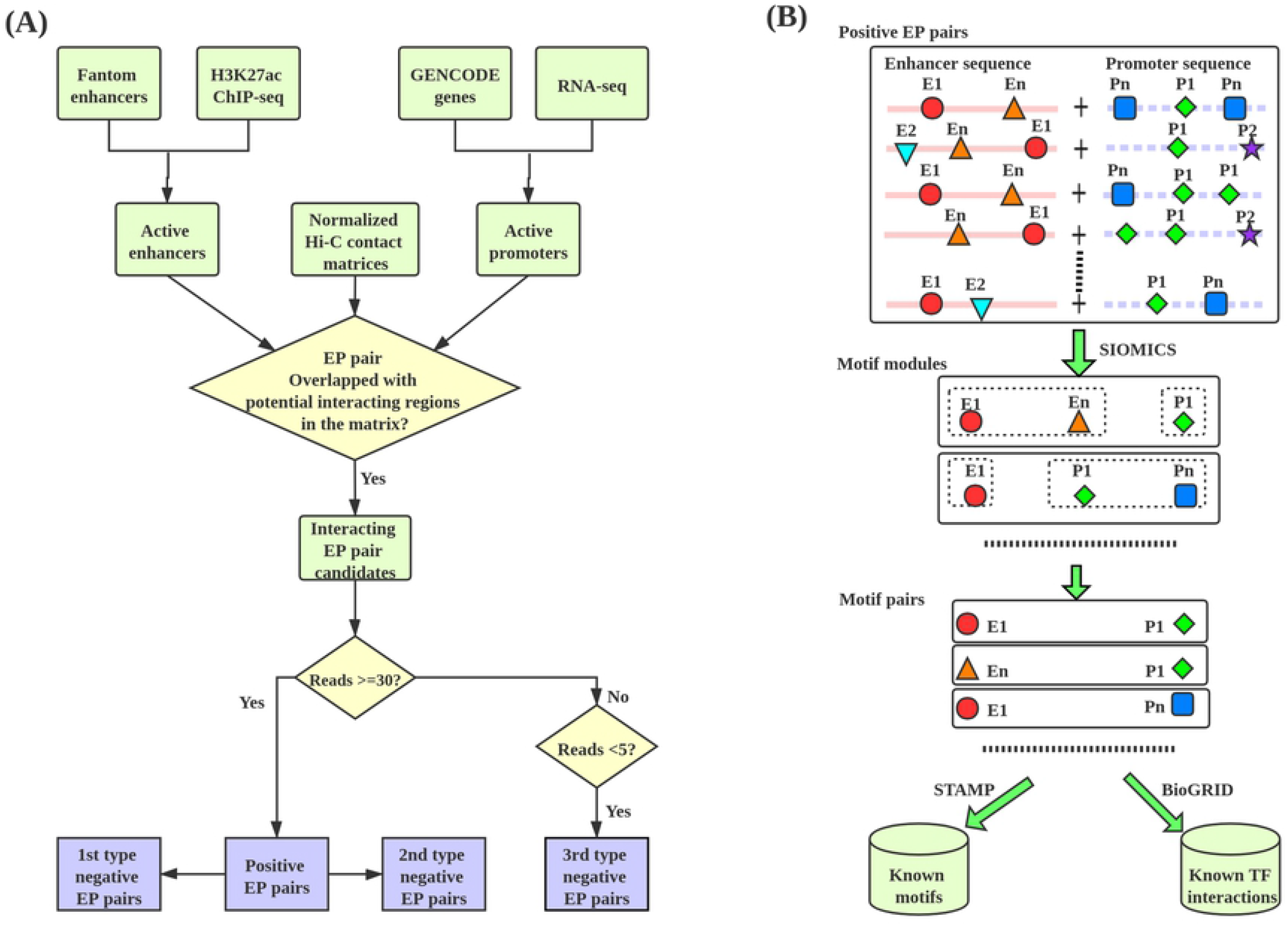
(A). The procedure to obtain positive and negative EP pairs. (B). The pipeline to study motif pairs in positive EP pairs.

To assess how well the predicted motif pairs facilitate the identification of true interacting EP pairs, we generated three types of negative EP pairs (Fig. 1A and Supplementary S1). The first type was the permuted version of the positive ones, in which the sequence of the enhancer in a negative EP pair was generated by randomly permuting the sequence of the enhancer in the corresponding positive EP pair, and the sequence of the promoter in this negative pair was from the random arrangement of the nucleotides in the promoter sequence of the corresponding positive pair. The second type of negative EP pairs was generated with randomly chosen genomic regions with the same lengths and a similar distance distribution to promoters as the corresponding enhancers as negative enhancers and the corresponding promoters in positive EP pairs as negative promoters. The third type was defined from the normalized Hi-C contact matrices with the cutoff 5, similar to the positive EP pairs (Fig. 1A). In brief, if a pair of genomic regions had fewer than 5 supported Hi-C reads, we called this pair of regions a negative pair of genomic regions. We then overlapped the negative pairs of genomic regions with the active FANTOM enhancers and active GENCODE promoters to obtain negative EP pairs. The first two types of negative EP pairs were used to assess whether the predicted motif pairs could distinguish non-EP pairs from positive EP pairs, while the third type was used to see whether they could separate the positive pairs from negative pairs. Similar to the positive EP pairs, the three types of negative EP pairs may contain false negatives and false positives. However, the proportion of such false positives was likely to be small.

### Non-redundant known motifs

We collected known TF binding motifs from JASPAR and CIS-BP databases [33, 37]. We compared every pair of motifs from these two sources with the tool STAMP [38]. As previously [39, 40], if two motifs had a STAMP similarity E-value smaller than 1E-05, we claimed they were similar. We obtained 649 non-redundant known motifs from the two sources by keeping only one motif in each group of similar motifs (http://www.cs.ucf.edu/~xiaoman/ET/EPmotif/).

### Discovery of motif pairs

To study motif pairs that may facilitate EP interactions, we obtained the DNA sequence of the enhancer and the promoter in every positive EP pair in each cell line. We then concatenated an enhancer sequence with its corresponding promoter sequence, if this enhancer and this promoter formed a positive EP pair (Fig. 1B). The obtained sequences were repeat masked (http://www.repeatmasker.org/papers.html).

Next, we applied the tool SIOMICS [39, 41] to these repeat-free concatenated sequences in every cell line to identify motif modules. A motif module is a group of motifs whose binding sites significantly co-occur in input sequences. Biologically, a motif module mimics the group of motifs for one TF and its cofactor TFs, where this TF and its cofactor TFs jointly bind to sequences to regulate a group of genes. We applied SIOMICS here because it considers multiple co-occurring sequence patterns to identify motif modules and motifs, which significantly reduces false positive predictions compared with the strategy to predict individual TF motifs separately. Moreover, it can de novo predict motifs, which thus does not depend on the limited number of known motifs available. It can also predict cofactor motifs for any variable number of cofactor motifs and has been shown good performance, which thus does not rely on prior knowledge of the groups of cofactor motifs. We ran SIOMICS on the repeat-masked sequences in each cell line separately, with the default parameters except *s* = 30 and *n* = 1500, which meant that we intended to identify up to 1500 motifs in a cell line and all motifs in a motif module must cooccur in at least 30 input sequences. The choice of 1500 is because there are about 1500 sequencing-specific binding TFs in the human genome [42]. SIOMICS assesses the statistical significance of the co-occurrence of every group of motifs by the Poisson clumping heuristic [43] and outputs the significant motif groups as motif modules (corrected p-value<0.01).

We then compared the predicted motifs in motif modules with the above non-redundant known motifs. A predicted motif was similar to a known motif if the STAMP E-value smaller than 1E-5. The TF(s) corresponding to this known motif was considered the TF(s) bound to this predicted motif. We also compared motifs predicted in different cell lines and claimed that two predicted motifs were similar if their STAMP E-value was smaller than 1E-8. The cutoff 1E-8 was used to compare the predicted motifs in different cell lines because motifs from the same source may be naturally more similar than motifs from different sources [44, 45].

Alternatively, we studied motif modules with known TF motifs. With the above 649 known motifs, we scanned the EP sequences with FIMO [46] in every cell line separately to obtain initial putative binding sites of known motifs. We then studied the co-occurring binding sites of groups of motifs by ChIPModule [47]. ChIPModule is similar to SIOMICS except that it considers the co-occurrence of the binding sites of known motifs in input sequences to predict motif modules. The initial putative binding sites of individual motifs defined by FIMO are thus refined based on the predicted motif modules by ChIPModule to reduce false positive predictions [47].

Finally, with the predicted motif modules, we obtained all pairs of motifs in every motif module. We then kept the pairs with the binding sites of one motif occurring in promoters and the binding sites of the other motif occurring in enhancers of positive EP pairs that contain the binding sites of the corresponding motif module. In other words, we filtered pairs that co-occurred in only enhancers or only promoters. These remaining pairs were the final motif pairs we considered, the TFs of which may be likely to interact and contribute to the interaction of the EP pairs (Fig. 1B).

### Homogeneous motif pairs

We consider a motif pair composed of the same motif as a homogeneous motif pair if this motif significantly co-occurs in both enhancers and promoters of the positive EP pairs. We apply two approaches to measure the significance of such a co-occurrence of the same motif to identify homogeneous motif pairs. In a first way, assume there are *N* positive EP pairs, and *n* has such a motif in both enhancers and promoters (based on FIMO scan). Assume the average promoter and enhancer length is *l_1_* and *l_2_* in this cell line, respectively. Also, assume this motif occurs *x* times in these *N*EP pairs. We calculate the p-value as *pbinom(n, N, p)*, where 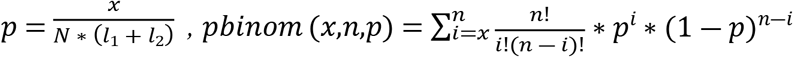. If this p-value is smaller than 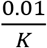 where *K* is the number of the predicted motifs in this cell line, we claim that this motif forms a homogeneous motif pair. In a second way, assume this motif occurs in *x* of the *N* enhancers and *y* of the *N* promoters based on the FIMO scan. We calculate the p-value with the same formula but different 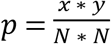. If this p-value is smaller than 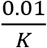 we claim this motif is significant. The difference between the two p-value calculations is that the first way considers the motif occurrence frequency and the sequence length difference, while the second way only considers the binary occurrence of a motif in a sequence and not the sequence length.

### Enrichment analysis of the predicted EP motif pairs

We compared the predicted EP motif pairs with known motif pairs of interacting TFs. We collected known directly and indirectly interacting TF pairs from the BioGRID database [48]. The direct TF interactions meant that two TFs physically interacted with each other. The indirect ones referred to pairs of TFs without direct interaction but directly interacting with a common third protein. There were 6820 pairs of direct and 120,277 pairs of indirect TF interactions in BioGRID, which involved 1520 and 1207 TFs, respectively. We then assessed the statistical significance of the enrichment of the motif pairs of known interacting TFs in the predicted motif pairs in every cell line by hypergeometric testing. In brief, assume there are N TFs and M pairs of TFs in BioGrid, among which there were *m* pairs that involved *n* TFs in the predicted EP motif pairs in a cell line. We calculate the hypergeometric testing p-value of enrichment of motif pairs of known interacting TFs 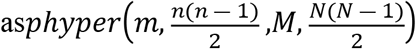, where 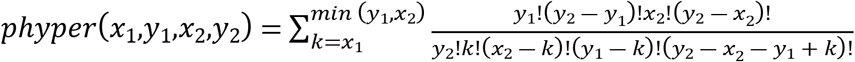. Also, we compare the predicted EP motif pairs with those predicted in a previous study [12]. This study predicted 298 pairs of TF interactions involved 61 TFs in GM12878, and 46 pairs of TF interactions involved 22 TFs in K562.

### Enhancer and promoter enriched motifs

Similar to the analysis of homogenous motif pairs above, we studied whether a predicted motif preferred to occur in enhancers or promoters by the binomial testing in two ways. One was to consider whether a motif occurred in significantly more enhancers or promoters. The other was to evaluate whether a motif appeared significantly more frequently in enhancers or promoters. The difference was the former emphasized the number of sequences and did not consider the length of sequences. In contrast, the latter considered both and emphasized the density of motif binding sites.

### Machine learning methods to distinguish positive from negative EP pairs

We attempted to address the following problem: given the positive and negative EP pairs, how well can we distinguish the positives from the negatives with the predicted motif pairs? We described each EP pair with a *4n+1* vector, where 4 entries were for each of the n motif pairs and one entry was for this pair’s positive or negative status. The four entries for a motif pair were the occurrence number of its motifs (based on FIMO) in the enhancer and promoter, respectively.

We applied four different methods to see how well these motif pairs distinguished positive EP pairs from negative EP pairs. The four methods were random forests, least absolute shrinkage and selection operator (lasso), decision tree and support vector machines [49–52]. We did 10-fold cross-validation to measure the performance of different methods. The four methods had similar F1 scores in separating positives from negatives. Because lasso utilizes fewer motif pairs while achieved similar performance, we presented our study with lasso to select the predicted motif pairs.

## Results

### The predicted motif pairs were likely to be biologically meaningful

We identified hundreds of motif pairs in interacting EP pairs in seven cell lines (Table 1). For every motif pair, at least one motif occurred in enhancers, and the other motif occurred in promoters of significantly many interacting EP pairs. These motif pairs were from the predicted motif modules, each of which contained 2 to 5 motifs. As mentioned above, a motif module is a statically significant group of co-occurring motifs, which represents the motif combination of a TF and its cofactors [53]. The predicted motifs, motif pairs, motif modules, and other information are available at http://www.cs.ucf.edu/~xiaoman/ET/EPmotif/.

**Table 1.**
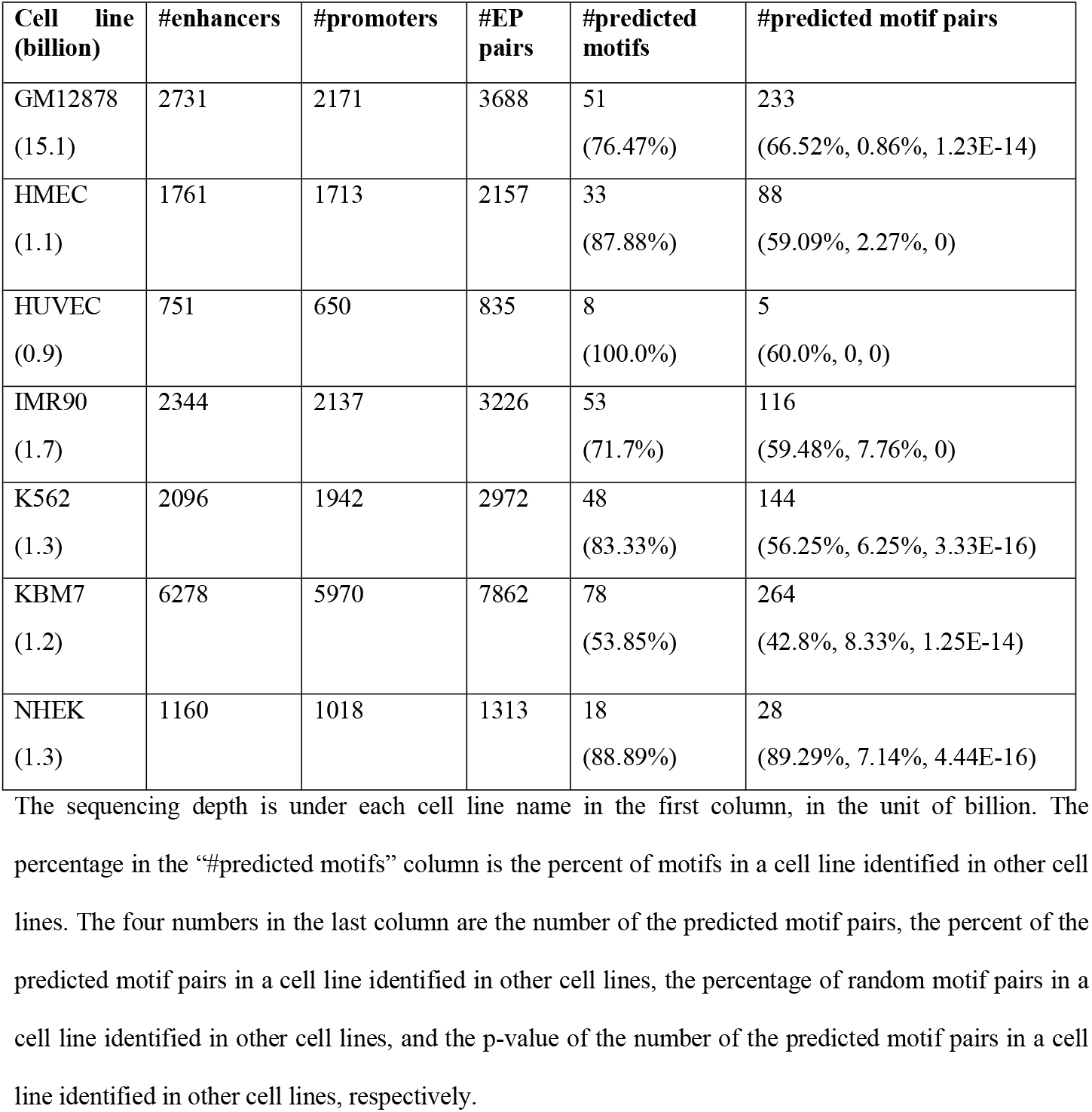
The predicted motif pairs in seven cell lines.

| **Cell line**<br>**(billion)** | **#enhancers** | **#promoters** | **#EP**<br>**pairs** | **#predicted**<br>**motifs** | **#predicted motif pairs** |
| --- | --- | --- | --- | --- | --- |
| GM12878<br>(15.1) | 2731 | 2171 | 3688 | 51<br>(76.47%) | 233<br>(66.52%, 0.86%, 1.23E-14) |
| HMEC<br>(1.1) | 1761 | 1713 | 2157 | 33<br>(87.88%) | 88<br>(59.09%, 2.27%, 0) |
| HUVEC<br>(0.9) | 751 | 650 | 835 | 8<br>(100.0%) | 5<br>(60.0%, 0, 0) |
| IMR90<br>(1.7) | 2344 | 2137 | 3226 | 53<br>(71.7%) | 116<br>(59.48%, 7.76%, 0) |
| K562<br>(1.3) | 2096 | 1942 | 2972 | 48<br>(83.33%) | 144<br>(56.25%, 6.25%, 3.33E-16) |
| KBM7<br>(1.2) | 6278 | 5970 | 7862 | 78<br>(53.85%) | 264<br>(42.8%, 8.33%, 1.25E-14) |
| NHEK<br>(1.3) | 1160 | 1018 | 1313 | 18<br>(88.89%) | 28<br>(89.29%, 7.14%, 4.44E-16) |

The sequencing depth is under each cell line name in the first column, in the unit of billion. The percentage in the “#predicted motifs” column is the percent of motifs in a cell line identified in other cell lines. The four numbers in the last column are the number of the predicted motif pairs, the percent of the predicted motif pairs in a cell line identified in other cell lines, the percentage of random motif pairs in a cell line identified in other cell lines, and the p-value of the number of the predicted motif pairs in a cell line identified in other cell lines, respectively.

The identified motif pairs were likely to be biologically meaningful, because we did not discover any motif pair when we carried out the same procedure in random sequences (the first type of negative EP pairs). We generated the corresponding number of random sequences as the original input for each of the seven cell lines by randomly permuting the nucleotides in each original sequence. We could not identify any motif in any cell line by applying the same procedure to these random sequences in each cell line. We thus could not identify any motif pair, implying the biological significance of the identified motif pairs.

The predicted motifs also corroborated the biological significance of the identified motif pairs. Motif pairs were composed of pairs of motifs predicted in the corresponding cell line. We noticed that we often independently identified the same predicted motifs across cell lines. On average, we independently discovered, more than 80% of the predicted motifs in different cell lines. The re-discovered motifs in multiple cell lines were not due to the shared EP pairs. We removed the shared EP pairs between every pair of GM12878, IMR90 and KBM7, which had the largest number of EP pairs, we could still find about 75% of the predicted motifs shared between every pair of the three cell lines. The independent discovery of the majority of motifs in other cell lines supported that these motifs were likely biological meaningful, which corroborated the function of the predicted motif pairs. Moreover, we also noticed that, on average, more than 55% of motifs in a cell line were similar to the annotated known motifs [33], further supporting the biological significance of the identified motif pairs in different cell lines.

The conservation of the identified motif pairs supported their biological significance as well. On average, more than 62% of motif pairs in one cell line were independently identified in other cell lines (Table 1). By randomly choosing the same number of motif pairs in each of the seven cell lines, we never had more than 10% random motif pairs discovered in other cell lines (p-value<1.25E-14). After removing similar motif pairs (both pairs of motifs had STAMP E-value<1E-08), we obtained 423 non-redundant motif pairs in seven cell lines. The conservation of the identified motif pairs suggests that these motif pairs were likely to be biologically meaningful.

### The predicted motif pairs were enriched with motif pairs of interacting TFs

In addition to the above evidence that supported the predicted motif pairs, we noticed that the TFs binding to these motif pairs is likely to interact. We obtained the TFs that may bind to a motif by comparing the predicted motifs with known motifs. In this way, we obtained the predicted TF pairs for the corresponding predicted motif pairs. We then compared the predicted TF pairs with the known interacting TF pairs extracted from BioGRID [48] (Material and Methods). We found that the predicted motif pairs were significantly enriched with those of interacting TFs in BioGRID.

In brief, in every cell line, we obtained TF pairs corresponding to the predicted motif pairs. Multiple TFs may bind to the same motif. We thus considered the TFs for a predicted motif in two scenarios: one was to include all TFs with their motifs similar to a predicted motif as the TFs of this predicted motif, and the other was to consider only the TF with the most similar motif as the TF of a predicted motif (STAMP E-value <1E-05 in both cases). In this way, we obtained two sets of TF pairs for the predicted motif pairs in every cell line (Fig. 2 and Supplementary S2). We then compared each of the two sets of TF pairs with the interacting TF pairs in BioGRID. The interacting TF pairs in BioGRID interacted directly or indirectly through a third common protein (Material and Methods). We found that the predicted interacting TF pairs were significantly enriched with the known interacting TF pairs in BioGRID in almost every cell line by hypergeometric testing (Fig. 2 and Supplementary S2).

**Fig. 2.**
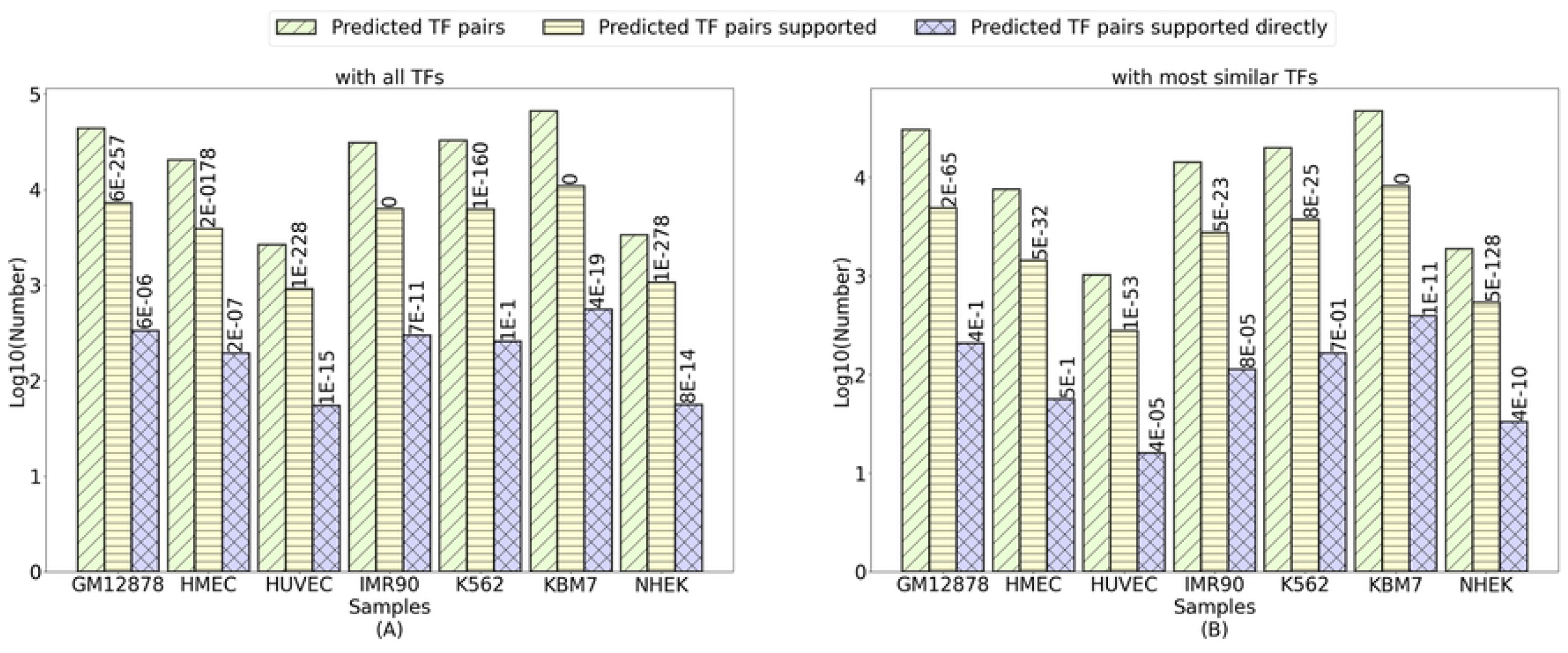
The predicted motif pairs are enriched with known interacting TF pairs.

Previously, Zhang et al. studied the ChIP-seq data and Hi-C data and computationally predicted the interactions of 61 TFs in GM12878 and 22 TFs in K562. We compared the predicted TF pairs in this study with theirs. There were 27 and 10 TFs in GM12878 and K562, respectively, shared by Zhang et al.’s study and this study. The two studies did not share all TFs, because certain TFs do not have sequence-specific binding motifs or do not have a known motif. There were 55 interactions in GM12878 and 4 interactions in K562 involving these shared TFs identified in Zhang et al.’s study. We identified 46 of the 55 interactions in GM12878 and 4 of 4 interactions in K562 (p-value 4.0E-27 and 0, respectively).

We investigated why we did not predict the remaining 9 interactions in GM12878. We found that we predict at least 8 of these 9 TF interactions. The motif pairs corresponding to these 8 TF pairs did not satisfy the motif similarity cutoff when we compared the predicted motifs with the known motifs with the STAMP E-value cutoff 1E-05. We also examined the motif pairs that were composed of known motifs and predicted in GM12878. We could identify all of the 55 TF pairs in GM12878, including all TF pairs of the missing 9 TF pairs. Also, we compared the TF interactions predicted by Zhang et al. with the BioGRID similarly. There were much fewer interactions predicted by Zhang et al. The p-values of the enriched TF interactions in their predictions were much smaller (Supplementary S3).

### The predicted motif pairs can help to distinguish positive EP pairs from negative ones

The above analyses suggested that the predicted motif pairs were likely to be biologically meaningful and may contribute to the interaction of EP pairs. We thus tested whether the predicted motif pairs could distinguish positive EP pairs from negative ones (Material and Methods). We found that the predicted motif pairs separated the positive EP pairs from the first two types of negative EP pairs well and reasonably distinguish the positive EP pairs from the third type of negative EP pairs (Table 2). All had the F1 score larger than 0.66.

**Table 2.** The accuracy of motif pairs in distinguishing positive EP pairs from three types of negative EP pairs based on lasso.

| Cell line | 1st type | 2nd type | 3rd type | #selected motif pairs | %selected motif pairs shared |
| --- | --- | --- | --- | --- | --- |
| GM12878 | (0.91,0.92,0.92) | (0.86,0.76,0.80) | (0.69,0.87,0.77) | (78, 96, 70) | 43/70=61.43% |
| HMEC | (0.90,0.90,0.90) | (0.85,0.72,0.78) | (0.52,0.99,0.68) | (66, 58, 36) | 26/36=72.22% |
| HUVEC | (0.83,0.88,0.85) | (0.67,0.79,0.70) | (0.51,1.00,0.67) | (5, 5, 5) | 5/5=100.00% |
| IMR90 | (0.91,0.92,0.92) | (0.91,0.79,0.84) | (0.50,0.99,0.67) | (56, 86, 43) | 18/43=41.86% |
| K562 | (0.91,0.90,0.91) | (0.87,0.75,0.81) | (0.50,0.97,0.66) | (71, 102, 53) | 25/53=47.17% |
| KBM7 | (0.91,0.89,0.9) | (0.90,0.72,0.80) | (0.59,0.90,0.71) | (107, 108, 53) | 16/17=94.12% |
| NHEK | (0.89,0.90,0.89) | (0.65,0.78,0.70) | (0.51,0.99,0.67) | (23, 24, 17) | 16/40=40.00% |
| Average | (0.89,0.90,0.90) | (0.82,0.76,0.78) | (0.55,0.96,0.69) | (58,68,40) | 55.46% |

The three numbers from the 2^nd^ column to the 4^th^ column are the precision, recall and F1 score. The second last column is the number of motif pairs selected by lasso in distinguishing positive from negative EP pairs for the three types of negatives in order. The last column shows the percentage of the selected motif pairs based on the third type of negatives by lasso in a cell line also selected in other cell lines.

We tried to determine how well the identified motif pairs could differentiate the positive EP pairs from the first two types of negative ones (Material and Methods). These two types of negative ones were “false” enhancers with random sequences or random genomic regions paired with promoters. We found that the predicted motif pairs told the positive EP pairs apart from the first type of negative EP pairs with an average precision of 0.89, and a recall of 0.90 in individual cell lines in 10-fold cross validation. Similarly, on average, the predicted motif pairs distinguished the positive EP pairs from the second type of negative EP pairs with an average precision of 0.82, and a recall of 0.76 in the 10-fold cross-validation (Table 2). For instance, in IMR90, the precision was larger than 0.90, and the recall was larger than 0.79 for both types of negative EP pairs.

We also studied how well the predicted motif pairs separated the positive EP pairs from the third type of negative EP pairs. In the 10-fold cross validation, the precision in all cell lines was from 0.50 to 0.69, while the recall was from 0.87 to 1 (Table 2). The much-reduced precision was likely because the number of negative EP pairs was much larger than that of positive EP pairs (Supplementary S1). We also noticed that the F1 score was decreasing from the first type of negatives to the third type of negatives, suggesting that it was more difficult to distinguish positive EP pairs from the third type of negatives than that from the first type of negatives. However, the F1 score was still above 0.66, indicating that the predicted motif pairs could facilitate to distinguish the true EP interactions from the false ones. In total, lasso selected 5 to 70 motif pairs in a cell line, which corresponded to 147 non-redundant motif pairs (Table 2). There were 30 motif pairs selected independently in at least two different cell lines.

We studied whether the predicted motif pairs in one cell line could distinguish the positive EP pairs from the third type of negative EP pairs in another cell line. The identified motif pairs in one cell line have similar precision and recall to distinguish the positive EP pairs from the third type of negative EP pairs in every other cell line to the predicted motif pairs from the corresponding cell line. This suggested that a large proportion of the predicted motif pairs in one cell line were likely to be conserved in another cell line. In other words, the predicted motif pairs represented conserved mechanisms across cell lines. We noticed that different cell lines shared not only the majority of the predicted motif pairs (Table 1) but also the majority of selected motif pairs used to distinguish positive EP pairs from the third type of negative EP pairs (Table 2).

### The selected motif pairs are likely to contribute to EP interactions

We studied whether the selected motif pairs contribute to EP interactions. Starting from the above 147 selected motif pairs, we identified pairs of TFs with their motifs similar to the selected motif pairs (STAMP E-value<1E-5). We could identify TF pairs for 72 of the above 147 selected motif pairs and 19 of the 30 motif pairs selected in multiple cell lines. For 64 of the 72 selected motif pairs and 18 of the 19 selected conserved motif pairs, their corresponding pairs of TFs interact in BioGRID. For at least 45 of the 72 pairs and 14 of the 19 pairs are shown to contribute to EP interactions in literature, among which 40 of the 72 pairs and 14 of the 19 pairs are supported by both BioGRID and literature (Supplementary S4). We provided two examples of the TF pairs corresponding to these selected motif pairs in the following. The remaining motif pairs and their functional support can be found in Supplementary S4.

An example of a novel motif pair selected is for the TF pair GATA1-ZNF423. GATA1 is known to bind to distal regions and physically interacts with ZFPM1 in the beta-major globin promoter [54]. Similar to ZFPM1, ZNF521, a paralog of ZNF423 that shares 65% of homology with ZNF423, is known to have a functional NuRD sequence at the N-terminal [55, 56]. Moreover, ZNF521 modulates erythroid cell differentiation through direct binding with GATA1 [57]. It is thus evident that GATA1-ZNF423 interaction is likely to facilitate EP interaction, which may be through the GATA1 interaction with the NuRD sequence at the N-terminal of ZNF423.

Here is another novel motif pair that may facilitate EP interactions. This selected motif pair is for the TF pair EBF1-ZNF143. In vertebrates, the EBF1 is demonstrated to have the role of controlling the higher-order chromatin structure [58]. ZNF143 is known to preferentially occupy anchors of chromatin interactions connecting enhancers and promoters [59]. Moreover, EBF1, ZNF143, and RAD21 have a three-way interaction in GM12878 [58]. It is thus likely that the interaction of EBF1-ZNF143 may contribute to EP interactions [19].

## Discussion

We de novo identified 423 motif pairs in interacting EP pairs. These motif pairs were likely to be biologically meaningful because they were statistically significant, conserved across cell lines, enriched with motif pairs of known interacting TFs, and so on. We also demonstrated that the predicted motif pairs could help to distinguish positive EP pairs from negative ones. We provided the predicted motifs, motif pairs, and other related information about these motifs and motif pairs at http://www.cs.ucf.edu/~xiaoman/ET/EPmotif/.

We identified 1183 motif pairs in interacting EP pairs with known motifs as well (Supplementary S5 and http://www.cs.ucf.edu/~xiaoman/ET/EPmotif/). We found that most of the identified motif pairs based on known motifs were similar to those de novo predicted ones in the corresponding cell lines. For instance, in KBM7, 94% of the identified motif pairs based on known motifs were similar to the de novo predicted motif pairs. A small fraction of the motif pairs based on known motifs were not discovered in the de novo predicted motif pairs, likely due to the STAMP E-value cutoff 1E-05 used.

We noticed that more than 55% of predicted motifs were similar to known motifs in one cell line. We also observed that more than 80% of the predicted motifs in one cell line were usually identified in other cell lines. In addition, we studied whether the predicted motifs preferred to occur in enhancers and promoters (Supplementary S6). Without considering the sequence length difference between enhancers and promoters, almost all motifs preferred to occur in promoters in all cell lines. When we considered the sequence length difference, where on average the promoters were three times longer than the enhancers, there was barely any motif preferring promoters to enhancers. Therefore, the majority of motifs occurred in both enhancers and promoters, with more frequent occurrence of their binding sites in enhancers.

We also checked whether there were homogeneous motif pairs that have the same motifs significantly occurring in both enhancers and promoters, such as the aforementioned CTCF-CTCF motif pair and the YY1-YY1 motif pair (Material and Methods). If we considered the sequence lengths, 78.6% to 93.1% motifs could form homogenous motif pairs that significantly co-occurred in positive EP pairs, including the CTCF-CTCF motif pair in six of the seven cell lines and the YY1-YY1 motif pairs in five of the seven cell line. Even if we did not consider the sequence length, we still could identify 13, 5, and 158 motifs that could form homogeneous motif pairs in GM12878, HMEC and KBM7. In this case, CTCF was still found in HMEC. If we lower the STAMP E-value cutoff when comparing the predicted motifs with known motifs, the predicted motifs similar to CTCF and YY1 were found in both GM12878 and KBM7. We provided two lists of homogeneous motif pairs based on the two different considerations at http://www.cs.ucf.edu/~xiaoman/ET/EPmotif/ for future validation studies.

Several directions may help to understand EP motif pairs better. First, although the identified motif pairs are likely to be useful in predicting EP interactions, they should be integrated with other features used previously [17, 19, 24] to fulfill their potentials. Second, a more comprehensive collection of enhancers and their condition-specific activity may improve the quality of the predicted motif pairs. The number of enhancers we used is relatively small compared with the collected enhancers in other resources [60, 61]. Third, with more annotated known motifs, it may be better to discover motif pairs directly from known motifs. We look forward to further exploring the EP motifs and their contribution to the interaction of the EP pairs.

## Acknowledgements

H.H. and X.L. designed the study. S.W., H.H. and X.L. analyzed data and wrote the manuscript.

## Supporting information

**Supplementary S1. Basic information about the data used in the paper.**

**Supplementary S2. Comparison of the predicted TF interaction with known ones in BioGRID.** In each entry, the information in order is the result based on all TFs for each predicted motif with STAMP cutoff 1E-05, and the result based on the most similar TF for each predicted motif with STAMP cutoff 1E-05. The p-value in 4^th^ and 6^th^ column is calculated based on hypergeometric testing. There are 1520 TFs in BioGRID, which are supported by GO. And there are 6820 TF pairs in BioGRID based on the 1520 TFs.

**Supplementary S3. Comparison between Zhang et al.’s study with BioGRID**

**Supplementary S4. TF pairs for 72 selected motif pairs by LASSO**

**Supplementary S5. Predicted motif pairs from known motifs.** The percentage in the “#predicted motifs” column is the percent of motifs in a cell line identified in other cell lines. The four numbers in the last column are the number of the predicted motif pairs, the percent of the predicted motif pairs in a cell line identified in other cell lines, the percentage of random motif pairs in a cell line identified in other cell lines, and the p-value of the number of the predicted motif pairs in a cell line identified in other cell lines, and the percentage of motif pairs found in the de novo predicted motif pairs independent of known motifs in the paper, respectively.

**Supplementary S6. Almost all motifs (SIOMICS) are likely to occur in both enhancers and promoters.**

